# Revisiting the high-dimensional geometry of population responses in visual cortex

**DOI:** 10.1101/2024.02.16.580726

**Authors:** Dean A. Pospisil, Jonathan W. Pillow

## Abstract

Recent advances in large-scale recording technology have spurred exciting new inquiries into the high-dimensional geometry of the neural code. However, characterizing this geometry from noisy neural responses, particularly in datasets with more neurons than trials, poses major statistical challenges. We address this problem by developing new tools for the accurate estimation of high-dimensional signal geometry. We apply these tools to investigate the geometry of representations in mouse primary visual cortex. Previous work has argued that these representations exhibit a power law, in which the *n*’th principal component falls off as 1*/n*. Here we show that response geometry in V1 is better described by a broken power law, in which two different exponents govern the falloff of early and late modes of population activity. Our analysis reveals that later modes decay more rapidly than previously suggested, resulting in a substantially larger fraction of signal variance contained in the early modes of population activity. We examined the signal representations of the early population modes and found them to have higher fidelity than even the most reliable neurons. Intriguingly there are many population modes not captured by classic models of primary visual cortex indicating there is highly redundant yet poorly characterized tuning across neurons. Furthermore, inhibitory neurons tend to co-activate in response to stimuli that drive the early modes consistent with a role in sharpening population level tuning. Overall, our novel and broadly applicable approach overturns prior results and reveals striking structure in a population sensory representation.

**Significance Statement:** The nervous system encodes the visual environment across millions of neurons. Such high-dimensional signals are difficult to estimate—and consequently—to characterize. We address this challenge with a novel statistical method that revises past conceptions of the complexity of encoding in primary visual cortex. We discover population encoding is dominated by approximately ten features while additional features account for much less of the representation than previously thought. Many dominant features are not explained by classic models indicating highly redundant encoding of poorly characterized nonlinear image features. Interestingly, inhibitory neurons respond in unison to dominant features consistent with a role in sharpening population representation. Overall, we discover striking properties of population visual representation with novel, broadly applicable, statistical tools.

---

**E**ach patch of the visual field is represented by a large population of neurons in primary visual cortex. This “population code” supports the performance of a huge diversity of downstream visual tasks, making it a topic of intense general interest in visual neuroscience. An important open question about neural population codes in V1 and beyond is their geometry (1, 2). Specifically, how does a population of *n* neurons make use of its *n*-dimensional activity space for representing external stimuli? One approach to this question is to examine the eigenvalues of the signal covariance matrix, which quantifies the correlations in a population’s (noiseless) responses over a collection of stimuli. If all eigenvalues are equal (corresponding to a “flat” eigenspectrum), the neurons are maximally uncorrelated, with each neuron encoding an orthogonal stimulus feature. Conversely, if the covariance contains only one non-zero eigenvalue, all neurons are perfectly correlated, meaning that they redundantly encode a single shared feature.

Recent work from Stringer *et al* 2019 (3) argued that an optimal population code must trade off competing demands of efficiency and smoothness. Efficiency, which relates to the code’s capacity for carrying information, requires a maximally flat eigenspectrum, so that the population takes full advantage of its dynamic range in all dimensions. Smoothness, on the other hand, relates to the property that nearby stimuli evoke nearby patterns of neural activity thus providing robustness to perturbations of stimuli and neural responses. Stringer *et al* argued that smoothness requires the eigenspectrum to decay at least as quickly as a power law with a slope of 1. Any slower decay of the eigenspectrum implies that the representation will not be smooth, so nearby stimuli elicit widely separated response patterns. Thus, the population code that maximizes efficiency while preserving smoothness is a power law with slope negative 1. Mathematically, the *i*’th eigenvalue of the (noiseless) response distribution should be *λi* = *ci*^−*α*^, where *α* = 1 is the power law exponent and *c* is a constant of proportionality.

To assess whether this property holds in mouse visual cortex, Stringer *et al* (3) introduced a novel method for estimating the signal eigenspectrum known as cross-validated PCA (cvPCA). On the basis of the cvPCA estimator applied to population responses in mouse primary visual cortex, they determined that the eigenspectrum both follows a power law and is at the critical limit of decay (*α* = 1). They interpreted this result as indicating that representations in V1 are as efficient as possible while maintaining smoothness.

Here we show that the cvPCA estimator provides a biased estimate of the signal eigenspectrum. We introduce a novel estimator for signal eigenspectra to overcome this bias. We then re-analyse the data from (3), and show that the signal eigenspectrum in mouse V1 systematically deviates from a power law. Rather, it is better explained by a broken power law, in which the largest eigenvalues follow a power law with shallow slope, and subsequent eigenvalues decay according to a different power law with steeper slope. Crucially, asymptotic decay of small eigenvalues under this model is not at the critical limit of *α* = 1, but decays significantly faster (∼20% steeper). We find that because of this form of the eigenspectrum population geometry is lower dimensional than previously thought and there are ten dominating eigenmodes that account for ∼30% of neural variation.

To gain insight into the population neural representations in mouse V1, we examined these dominant dimensions of the population response. We found that some dimensions often recapitulated classical selectivity for spatial frequency and orientation that has been reported in primary visual cortex (4, 5) but with far higher fidelity than single neurons. However, other dimensions, that were also robustly encoded, were unexplained by classic models indicating that difficult to characterize single neuron tuning (6, 7) is highly redundant across neurons. Furthermore, we found that inhibitory neurons’ contribution to these dominant dimensions tended to be larger and more uniform than the excitatory cells consistent with a role in sharpening population tuning analogous to single neuron level effects of inhibition (8). Overall, these findings highlight the importance of examining sensory representations at the population level to uncover emergent coding properties that are not apparent from single neuron responses alone.

## Results

Neural tuning refers to a neuron’s average or “noise-free” response for a collection of stimuli. ((9–11), Fig 1A). The “population” or “joint” tuning of a neural population is thus an *n*-dimensional cloud of points defined by the mean responses of all *n* neurons in the population over a particular stimulus set (Fig 1B). To quantify the geometry of this joint tuning, we can compute the eigenvalues of its covariance, which describe the variance of this cloud of points along each axis in a set of *n* orthogonal axes known as eigenmodes. This set of eigenvalues, sorted from greatest to smallest, is known as the signal eigen-spectrum. Estimating the signal eigenspectrum from neural population recordings is a challenging statistical problem. In high dimensional settings, the number of stimuli that can be shown in an experiment may be smaller than the number of neurons in the population. Moroever, neural responses are noisy, meaning that multiple presentations of each stimulus are required to accurately estimate the mean response to each stimulus (Fig 1E).

**Fig. 1.**
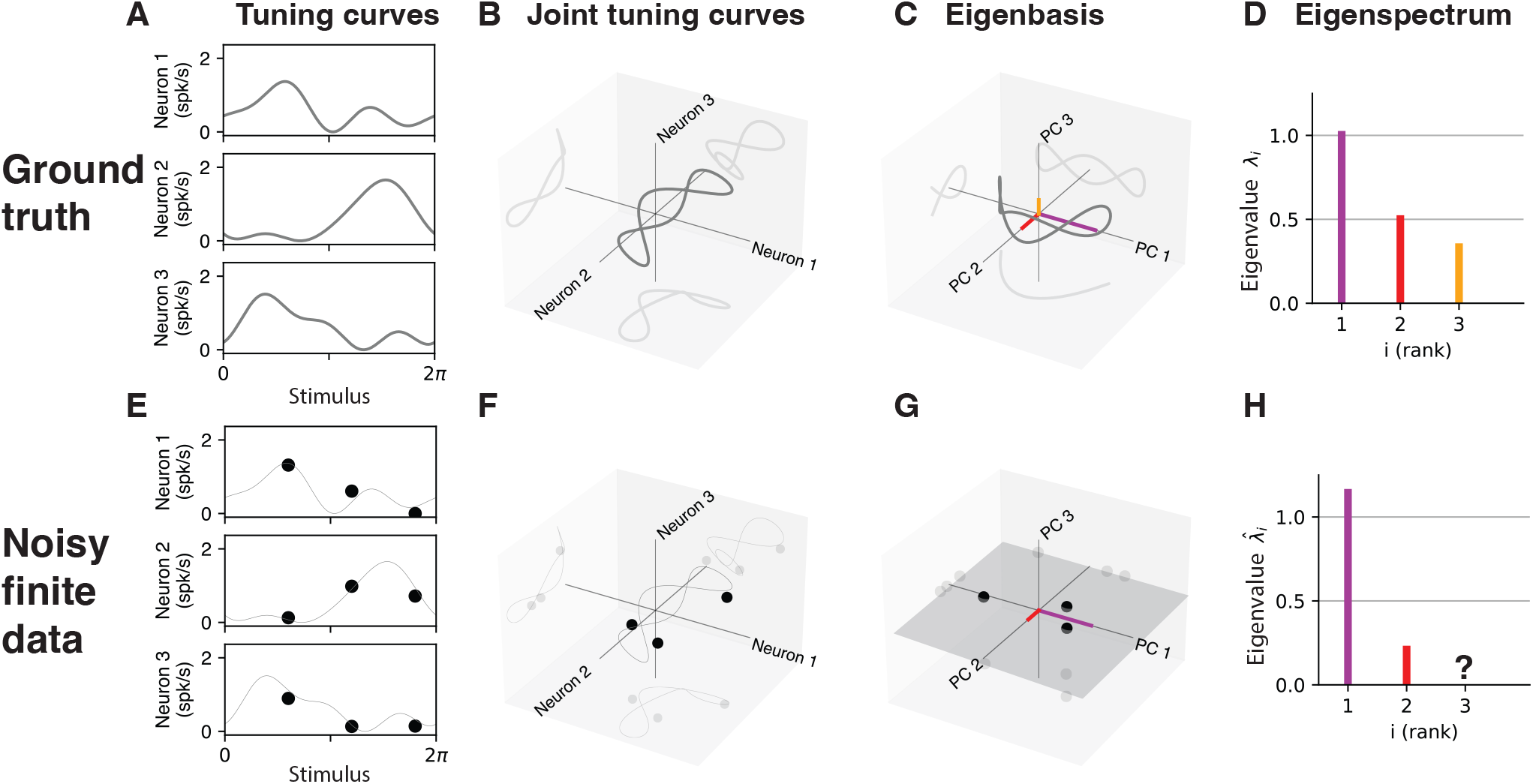
The signal eigenspectrum and the challenge of estimating it. **(A)** Tuning curves of three neurons for a 1-dimensional stimulus (gray traces). **(B)** Same three tuning curves plotted jointly in a 3D response space. **(C)** Joint tuning curve centered and plotted along principal axes of variation. **(D)** Eigenspectrum, which describes the variance along each principal component of the joint tuning curve. **(E)** Noisy estimates of individual tuning curves at the same three points along the tuning curve (black points). True tuning curve is unknown (light grey trace). **(F)** Noisy estimate of the joint tuning curve (black dots). **(G)** Estimated joint tuning curve centered and rotated to align with its principal components; the resulting curve is 2-dimensional, since 3 points defined a plane. **(H)** Eigenspectrum of the estimated joint tuning curve. Only two eigenvalues are non-zero, and thus later eigenvalue of true tuning curve are missing.

Principal components analysis (PCA) applied to trial averaged responses provides a standard method for estimating the signal eigenspectrum. It finds a sequence of orthogonal directions in neural response space that capture maximum response variance. However, this approach leads to two sources of bias (1) trial-to-trial noise covariance can corrupt estimates of the underlying signal covariance, (2) even in the absence of trial-to-trial noise, finite sampling of stimuli will bias estimates of the eigenspectrum— for example if there are fewer stimuli than neurons (*d < n*) then the sample covariance matrix will only have *d* non-zero eigenvalues, thus *n*−*d* eigenvalues. For example, three observations in neural response space (Fig 1F) that have been centered can always be described perfectly by two dimensions (Fig 1G) and thus eigenvalues with indices above 2 will be 0 (Fig 1H).

The cvPCA estimator was proposed as a solution for bias introduced by trial-to-trial noise. The estimator for the *i*th eigenvalue is formed by computing the estimated signal variance (using an unbiased estimate of the signal covariance) along the *i*th eigenvector of an unbiased estimate of the total covariance—signal plus noise covariance (see Methods, cvPCA). The noise covariance can then, for example, perturb the first eigenvector into a direction that is not the direction of maximal signal variance. Thus if the ordered signal and noise covariance matrix eigenvectors are not perfectly aligned, the cvPCA estimator will converge to the incorrect values (See Fig 2F). Additionally, cvPCA cannot estimate signal eigenvalues greater than the number of stimuli, which limits its ability to accurately recover lawful relationships in the decay of small eigenvalues, which is of fundamental importance for the dimensionality of neural populations.

**Fig. 2.**
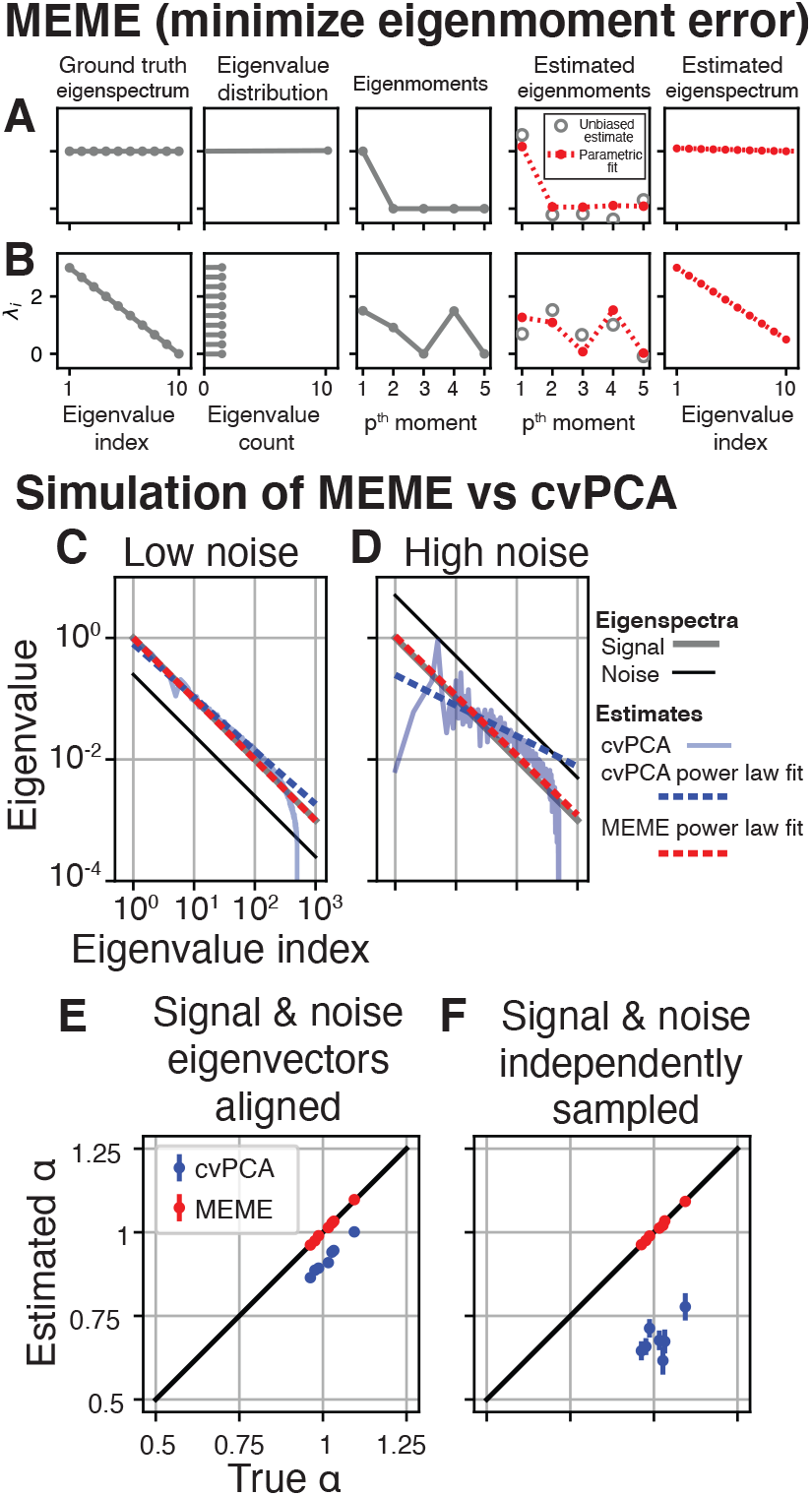
MEME estimator and validation in simulation. **(A)** Schematic of MEME method applied to uniform eigenspectrum **(B)** and to eigenspectrum with linear decay. (C) Comparison of MEME (red) and cvPCA (blue) estimates of power law in 1000 dimensions with low noise (signal eigenspectrum grey above noise eigenspectrum black) and high number of stimuli (m=500). In the case of cvPCA, a power law is estimated by fitting a line in log-log coordinates. We fit this line to eigenvalues along eigenvalues 2-50 (blue dotted) matching the proportions used in Stringer *et al*, (3) (D) Same simulation but with high noise. **(E)** Comparison of estimators on data draw matching the distribution of experimental data from Stringer *et al*, but where the signal and noise eigenvectors are the same and the eigenspectrum is set to be a power law matching the slope estimated by the cvPCA procedure. **(F)** Simulation where signal and noise eigenvectors are independently formed from noise.

To overcome these limitations, we introduce a novel method for estimating the signal eigenspectrum from noisy neural recordings by exploiting a recently developed estimator for the moments of the eigenvalue distribution.

### A moment-based estimator for the neural eigenspectrum

We developed a novel estimator that, up to a good approximation, does not suffer from any of the three biases we have described. We did so by finding unbiased estimates of signal ‘eigenmoments’, the *p*th moment being the signal covariance eigenvalues to the *p*th power averaged 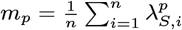, then finding the best fit eigenspectrum to these unbiased estimates. We found our signal eigenmoment estimator by extending the results of Li *et al* (2014) and Kong & Valiant (2017).

To provide intuition into this approach it is useful to consider the centered eigenmoments of two different eigenspectrum (Fig 2). If the eigenspectrum is flat (Fig 2A, column 1), implying each neuron’s tuning is mutually orthogonal to all other neurons’ tuning, then the distribution of eigenvalues will be a delta function centered at the average variance of the neurons (Fig 2A column 2). The first eigenmoment is the mean of the eigenvalues and thus is also equal to the average variance of the neurons but all other moments are zero because there is no spread to the distribution (Fig 2B column 3 traces go to zero after *p* = 1). If an unbiased estimate of the first eigenmoment was obtained, 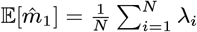, and we knew the eigenspectrum was flat we would have an unbiased estimator of the eigenspectrum 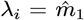. If we are unwilling to make such a strong assumption we can choose a more flexible parametric form of the eigenspectrum, for example it is linear as a function of the index, and fit it to unbiased estimates of higher order moments (Fig 2A column 4, error between red dashed parametric eigenmoments and grey open circle unbiased estimates is minimized) and the eigenspectrum associated with those eigenmoments serves as an estimate of the ground truth eigenspectrum (Fig 2A column 5, red dashed trace).

If the eigenspectrum is not flat but decreases linearly with the index (Fig 2B) then the distribution of eigenvalues will be uniform (Fig 2B column 2). The second eigenmoment will be non-zero because of this spread but every odd eigenmoment, but one, will be 0 because the distribution of eigenvalues is symmetric (Fig 2B column 3). More generally, the eigenspectrum is uniquely specified by its eigenmoments.

Critically the expected value of our estimates of signal eigenmoments are unbiased so do not depend on the rank of the data used to estimate them nor by corrupting noise regard-less of its covariance structure. Furthermore we prove these eigenmoment estimates are unbiased regardless of the data’s distribution, provided finite moments, thus these guarantees are broadly applicable. We now show that for typical ranges of parameters in neural data our estimator is highly accurate and overcomes issues with the prior estimator cvPCA.

### Validation of estimator in simulation

To demonstrate the key properties and effectiveness of our estimator we ran a simulation where both signal and noise eigenvalues followed a power law. cvPCA is the only other estimator that has been proposed to specifically estimate the signal eigenspectrum thus we compare our estimator to it.

We first simulated *d* = 1000 neurons, *m* = 500 stimuli, and *n* = 2 repeats. This corresponds for example to a typical calcium recording experiment to characterize sensory tuning in a population recording. To estimate a power law Stringer *et al* (3) fit a line in log-log coordinates only for eigenvalues with indices between 11-500 out of 10,000 eigenvalues. Here we matched this procedure for a smaller number of neurons with a scaling factor of 1,000/10,000 to fit eigenvalue indices between 2-50. In the case where noise was low (Fig 2C grey signal eigenspectrum above black noise eigenspectrum) the MEME estimator performs well (red overlap grey) and cvPCA performs similarly. When we increased the noise level (Fig 2D, black above grey) we found that early cvPCA estimates tended to dramatically mis-estimate the true signal eigenspectrum (transparent blue trace on left well below grey) and this led to mis-estimation of the power law fit to the cvPCA estimates (dashed blue traces do not align with grey). Whereas the MEME estimate continued to accurately estimate the form of the power law (red dashed trace overlaps grey). It is possible that for a different choice of range the cvPCA estimated power law could have been more accurate but it is unclear how to apriori choose this range when the true power law is not known. Thus in simulation we discovered biases in the approach of fitting a power law to cvPCA estimates. We now consider if these biases could have affected results in the original study of Stringer *et al*.

We found unbiased estimates of the signal and noise covariance of the original seven recordings of mouse primary visual cortex to natural images then enforced a true power law signal eigenspectrum that matched the slope estimated from cvPCA (see Methods, ‘Simulation procedure’). We then simulated data from this distribution and fit the signal power law using the original cvPCA approach and MEME. High dimensional signal and noise eigenvectors are difficult to estimate so we chose two extremal cases for our simulation. In the first case we aligned the signal and noise eigenvectors and found that cvPCA consistently under estimated the slope of the power law exponent *α* (Fig 2E blue points below black diagonal) whereas MEME accurately recovered the slope (red points overlaps black diagonal). We then ran the same simulation but where signal and noise were independently sampled and found an even larger downward bias of cvPCA while MEME remained accurate (Fig 2F). Thus we expect that regardless of the relationship between signal and noise the cvPCA power law exponent estimate is biased downwards but less so to the degree that signal and noise are aligned (see Methods, ‘cvPCA’). Signal and noise correlation are known to co-vary (12–16), but see (17), thus it is plausible that in neural data the bias of cvPCA may be ameliorated somewhat.

Given that cvPCA returned biased estimates of the signal eigenspectrum on simulated data matching the distribution of the original data, whereas MEME was accurate, we next examined whether estimates of signal eigenspectrum on the original data using the two different methods diverged.

### Application to estimation of power law eigenspectrum in mouse primary visual cortex

We re-analyzed the original data from Stringer *et al*, (3), responses from ∼10, 000 neurons across a patch of primary cortex (Fig 3A). Two repeated responses of all neurons to a set of ∼2, 800 stimuli were collected (Fig 3BC) and these responses were mean centered neuron-wise (for details of calcium response pre-processing see Methods). In general we found these neurons tended to respond to a restricted region of the stimuli (Fig 3D, average power of estimated linear receptive fields across all neurons of example recording). We first applied cvPCA to an example recording and fit a power law finding that it had a slope near 1 (Fig 2E blue, *α* = 0.96). When we fit a power law using MEME we found a significantly shallower slope (*α* = 0.90). Yet the eigenmoments of this MEME estimated power law systematically deviated from the unbiased estimates of the raw data’seigenmoments, implying that a power law was a poor fit to the data (Fig 3F pale red points deviate beyond CI’s of grey points). Similarly the eigenmoments of the cvPCA estimate did not match the data’s eigenmoments (blue points deviate beyond CI’s of grey points). This motivated us to consider more flexible eigenspectrum functions. Given that the original study formed predictions with respect to the exponent of a power law, we fit a piece-wise power law to obtain a more flexible model while still being able to make direct comparisons to their predictions. We found that in all cases, accounting for model degrees of freedom, the broken power law fit the eigenmoments of the data better than a power law (Fig 2F red dots within CI’s of grey, see supplementary information Fig S1 for statistical tests across all recordings). The broken power law had an initial shallow power law and a tail power law that was much steeper (Fig 3E red trace slope initial ∼0.5 then ∼1.2). This form of eigenspectrum was similar across all recordings (Fig 3G red traces overlap). The slope of signal eigenspectrum tail was consistently higher for the MEME than cvPCA estimates (Fig 3H, MEME average *α* = 1.20, cvPCA average *α* = 1.01). These findings are inconsistent with two claims from the original study. First, the eigenspectrum of population responses in mouse visual cortex is inconsistent with a power law, we find it is far better described as a broken power law. Second, at no point does the eigenspectrum decay at a critical rate near *α* = 1, instead it initially decays 50 % more slowly and then 20 % faster in the tail of the eigenspectrum.

**Fig. 3.**
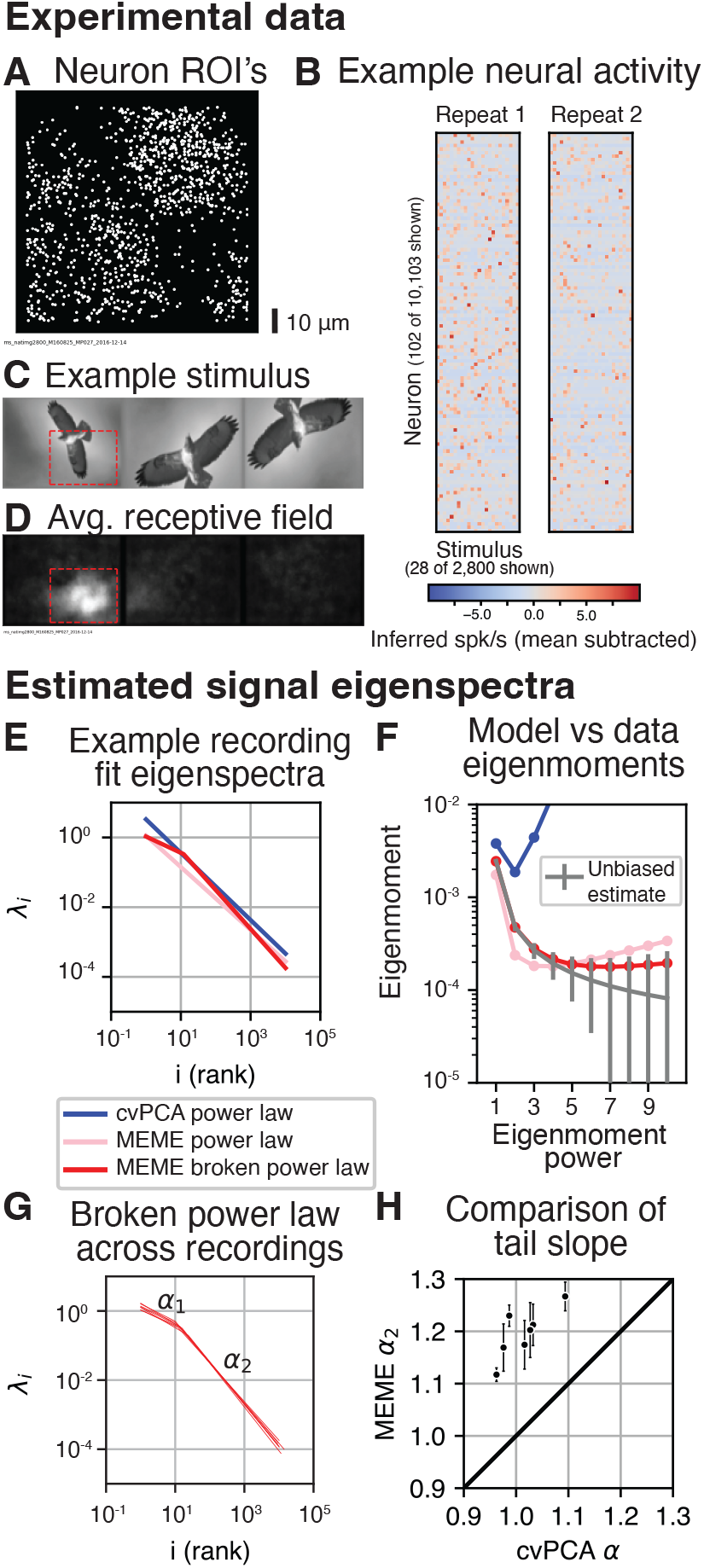
Fit of Stringer *et al*, (2019) data using cvPCA and MEME. **(A)** Positions of 1,011 neurons 10,103 recorded from primary visual cortex of mouse.**(B)** Example neural data, two repeats of the simultaneous responses of neurons to the same set of stimuli. **(C)** Example stimulus shown to mice drawn from ImageNet. **(D)** Estimate of the population receptive field (average power of estimated linear receptive fields of all neurons). Bounding box used for visualizing linear RFs (red dashed). **(E)** Signal eigenspectrum fit to neural data: a power law fit to cvPCA estimates following the methods of Stringer *et al*, (blue), a power law fit to unbiased estimates of signal eigenmoments pale red, and a broken power law fit to the same eigenmoments (red). **(F)** Unbiased estimates of eigenmoments with 95 % CIs compared to the eigenmoments corresponding to the eigenspectrum in (E). **(G)** Across all recordings (n=7) the best fit broken power law eigenspectrum (red). **(H)** The power law exponent estimated by cvPCA plotted against the exponent of the tail of the broken power law estimated by MEME (*α*_2_ see (G)). Plotted are individual estimates with 95 % CI’s for the MEME estimates and cvPCA (black points).

Despite the more rapid decay in the tail of the eigenspectrum the overall dimensionality (as quantified by the participation ratio (18, 19)) was on average 1.68 times higher than a power-law with a slope of 1 would predict. Thus the dimensionality of primary visual cortex is much higher than previously thought but because of only ∼10 dominating modes. The number of dimensions needed to capture 75% of the variance of population tuning, another metric of dimensionality (20), is actually lower (under the fit broken power law on average 357 eigenvectors are needed whereas for a power law with a slope of one 902 eigenvectors are needed). This contradiction in two metrics of dimensionality is precisely because the dimensionality increase quantified by the participation ratio is driven by the first ten modes and once these are accounted for the remaining variation is captured rapidly by successive dimensions (see supplementary information Fig S2).

These results imply there are two distinct regimes of joint encoding in mouse visual cortex. A high dimensional regime where ten dominating features of the stimulus have a similar magnitude of effect on the population and a low dimensional regime where the remaining variation of tuning is rapidly absorbed. This led us to examine the encoding properties of the dominant modes.

### Characterization of population tuning

The signal eigenspectrum corresponds to a decomposition of neural responses into directions of maximal signal variation across stimuli. We will call these directions of maximal variation in neural response space “neural eigenmodes” (identical to the eigenvectors of the neural signal covariance matrix) and the variation in the scale of these modes across stimuli “eigenmode tuning” (identical to the eigenvectors of the stimuli signal covariance matrix). We estimated these by respectively calculating the eigenvectors from unbiased estimates of the signal covariance over stimuli and neurons. The neural eigenmode loadings tended to be sparse with most weights near 0 but a few very large weights (Fig 4A black trace concentrated around 0). For the first mode we found a bias in the sign of the loadings (Fig 4A first row black trace biased upwards) with 69 % positive. Thus, the most variation in neural signal variation can be described as uniform excitation on a subset of neurons. To gain insight into the tuning of this eigenmode we fit a linear model that predicted eigenmode tuning from a linear combination of stimuli pixels (Fig 4B orange traces, *R*^2^ = 0.2). Visualizing the weights on stimuli pixels we found classic center surround tuning (Fig 4C). Thus, surprisingly, a substantial fraction of the variation in the first dominant mode of neural tuning could be explained by a classic model of early visual selectivity. To gain further insight we examined the stimuli that evoked the three highest and lowest responses of this eigenmode and compared them to the linear component of the responses (Fig 4D first row of black and orange outlined images). Qualitatively comparing the two sets we judged that the eigenmode tuning was driven by higher spatial frequency image structure than the linear component. Careful analysis of more flexible models could gain greater insight into the non-linear component of eigenmode tuning (i.e., the systematic prediction errors of the linear models). Examining the linear receptive fields of other recordings we repeatedly observed clear selectivity for spatial frequency matching the scale of the population receptive field, a diversity of orientation selectivity, and phases (Fig 4F left to right). Thus classical primary visual cortex receptive field properties drive a significant amount of variation in the top eigenmodes of mouse primary visual cortex.

**Fig. 4.**
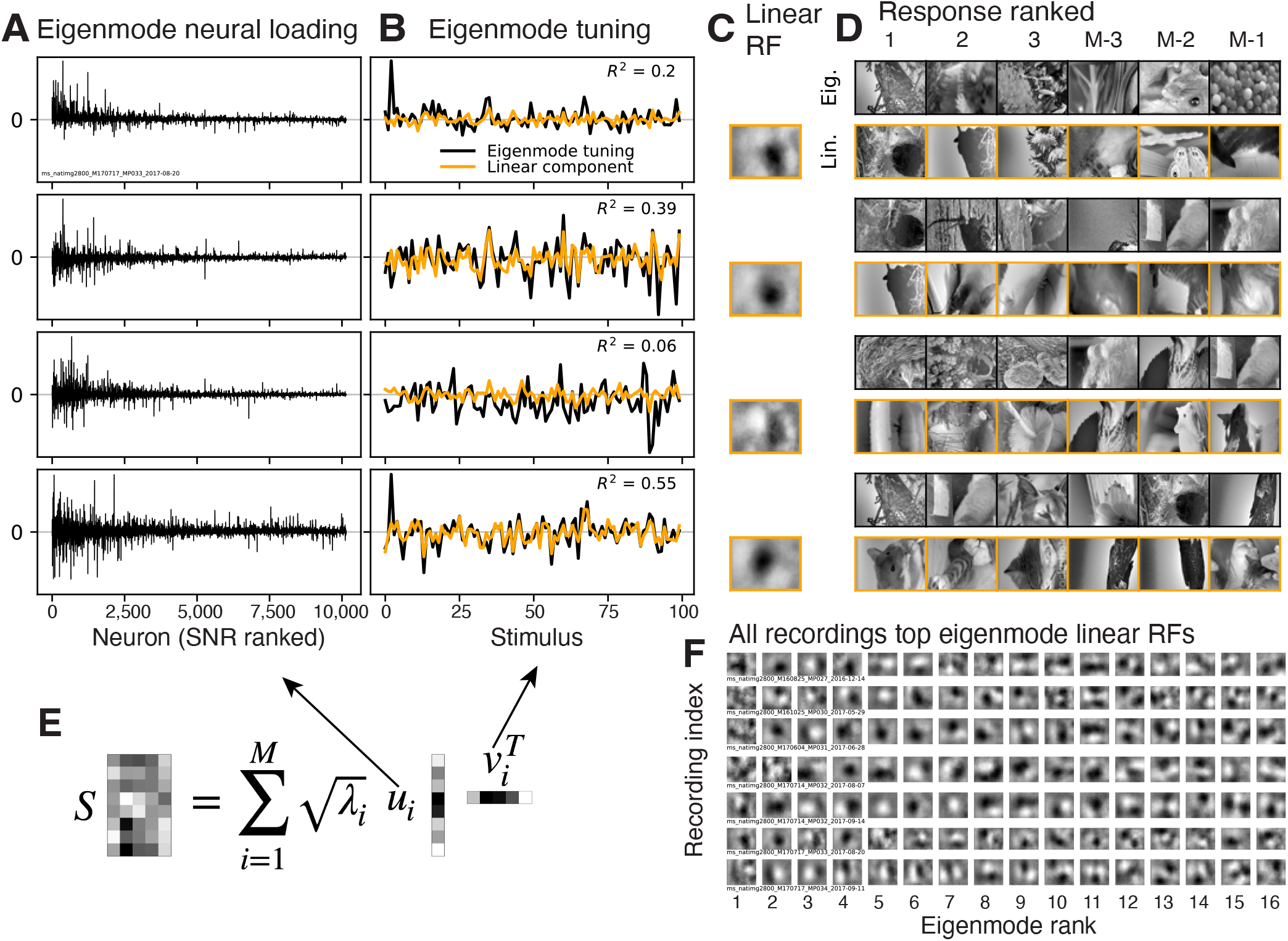
Analysis of population neural tuning. **(A)** Signal eigenmode loadings on each neuron (estimated left-singular vectors of noiseless responses, neuron by stimuli matrix) plotted against SNR of the neuron. **(B)** Signal eigenmode tuning (right-singular vectors) in black, least squares fit of image pixels to eigenmode tuning in orange. **(C)** Visualization of linear receptive field of eigenmode tuning (dot product of linear RF pixels with image pixels gives orange trace in (B)). **(D)** Stimuli that gave the top and bottom three responses from eigenmode tuning (black outlined top row) and the linear receptive field (orange outlined bottom row). **(E)** Formula for reconstruction of neural signal matrix (rows neurons, columns stimuli) from eigenmode neural loadings (left singular vectors of signal matrix which in the limit of infinite stimuli equals the eigenvectors of the signal covariance matrix) and tuning vectors (right singular vectors). **(F)** Linear receptive fields from all recordings of responses to natural images (rows) ranked by eigenmode (columns).

A normative explanation for the presence of signal correlation between sensory neurons is that it can improve the fidelity of the signals encoded in common across a population of neurons (21–23). Here we quantified the scale of this effect by measuring the noise corrected SNR (24) of eigenmodes and single neurons. We estimated eigenmode neural loadings with 2,000 stimuli then projected neural responses to the rest of the stimuli (∼ 300 −800) onto those loadings and calculated SNR across the two repeats. We found that tuning for early eigenmodes had higher fidelity than the average neuron (Fig 5A, grey trace above black dashed for indices 1-10). This was a consistent result across recordings with the first 10 eigenmodes having an average SNR at least 4.9 times greater than that of the average single neuron (average SNR 7.7 times greater). Furthermore, eigenmode SNR is likely underestimated because our estimates of signal eigenvectors are noisy. We can conclude that a hypothetical downstream region could more easily decode the feature encoded by an early eigenmode than a typical neuron because of the structure of signal correlations.

**Fig. 5.**
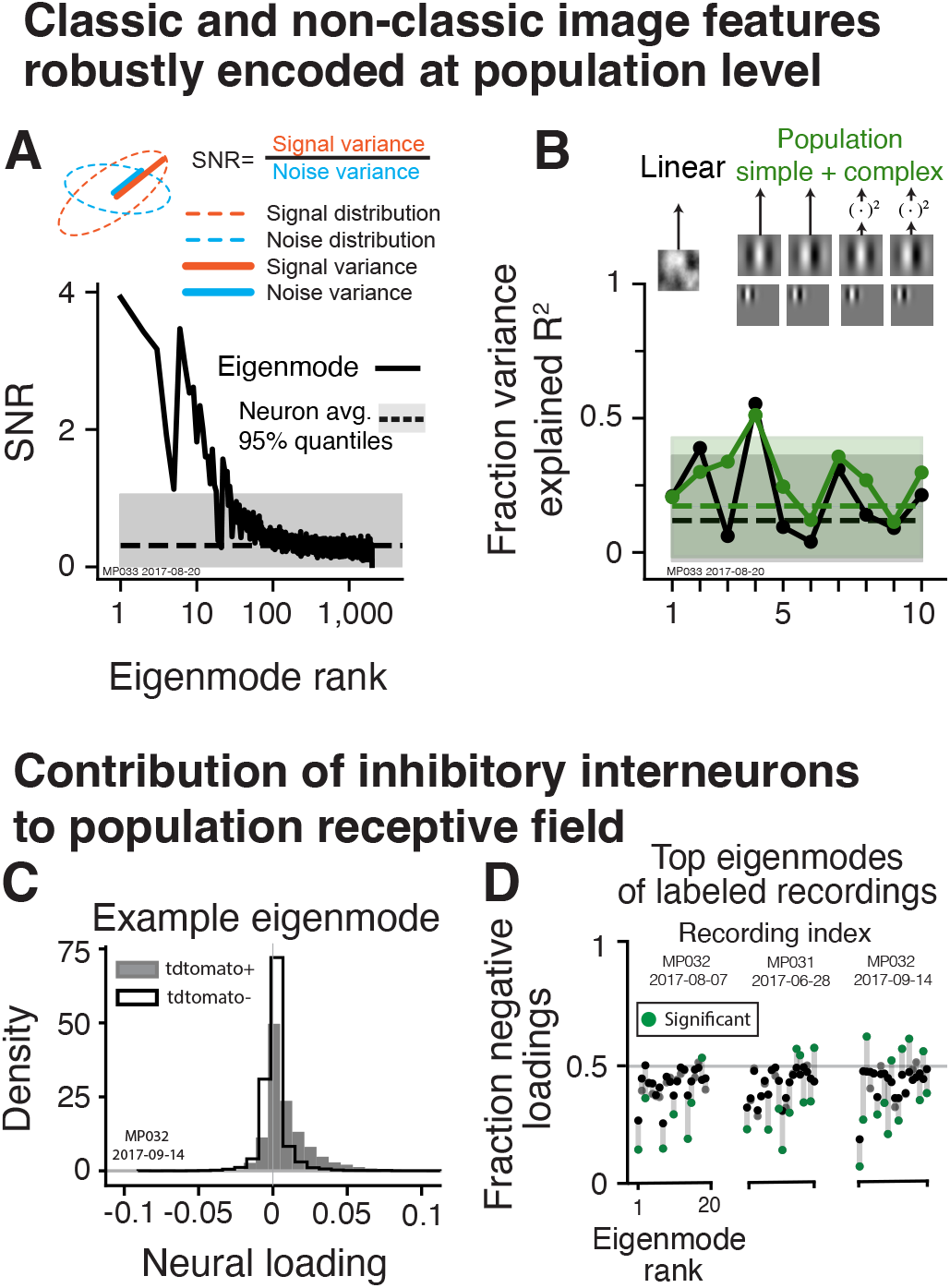
Population and single neuron tuning and the distinct contribution of inhibitory interneurons to the population receptive field. **(A)** Estimated SNR of ranked signal eigenmodes (black solid trace) compared to average SNR of individual neurons (black dashed horizontal line) along with 95 % quantile (transparent grey). **(B)** Fraction variance explained (corrected for noise and model degrees of freedom) by linear model (black) and simple and complex cell model multi-scale population (green) for single neurons on average and top ten eigenmodes.**(C)** The distribution of eigenmode neural loading on neurons identified as inhibitory (grey) and other neurons (black) for the 2nd eigenmode in example recording. **(D)** Across the top 20 eigenmodes for the three tdtomato+ labeled recordings the fraction negative loadings for inhibitory and other neurons (respectively gray and black), green dots indicate where these fractions are significantly different (p<0.001).

Ultimately these dominant eigenmodes, which we found robustly encode image features, are the result of redundancy in the tuning between individual neurons. We observed that often early eigenmodes possess receptive field properties typical of the classic characterization of individual neurons in primary visual cortex (Fig 4F). It might be expected that if the selectivity of all 10,000 neurons were restricted to the relatively small linear subspace spanned by a narrow band of spatial frequencies the top eigenmode tuning would inevitably recapitulate this structure. Yet, it is well known that classic models do not often predict the bulk of variation in single neuron tuning, in fact the tuning of individual neurons are notoriously difficult to predict across natural images even with flexible data driven models (6, 7). Indeed, when we estimated the ability of a linear filter to predict neural responses, by regressing the pixels of the images on the neural responses and estimating *R*^2^ using a noise corrected estimator (24) (see Methods, Estimation of model performance), we found that on average less than a quarter of neuronal signal variance could be predicted (Fig 5B dashed black trace). We also fit a basis of gabor filters and their squares (a multi-scale ensemble of classic simple and complex cell models (6, 25)) and found that on average predictive performance increased only slightly (green dashed above black). Thus a minority of single neuron tuning is characterized by linear or classic receptive field properties thus a majority is non-linear and not characterized by classic models. Unlike the stereotyped classical receptive fields it is not obvious whether or not this single neuron selectivity will be robustly represented at the population level. Each neuron’s unexplained tuning could be orthogonal. Yet, we find that it is often highly redundant at the population level: less than a quarter of the variation in the top eigenmode can be captured by a classic model (Fig 5A beginning of black and green solid trace). These eigenmodes are noisy estimates so it is not clear how much of their tuning is ‘explainable’ but we find that later, more difficult to estimate, eigenmodes can often be better explained (black trace peaks at eigenmode 4 with 50 % variance explained). Across recordings we find that it is typical for some modes to have up to half their variance explained while others, often the first mode, have less than a quarter explained (see supplementary information Fig S3). These results suggest there are distinct single neuron tuning properties that are highly redundantly encoded across primary visual cortex but that are not well characterized by classic models of primary visual cortex. Redundancy implies these tuning properties are of particular import to the organism and yet it remains unclear what these tuning properties are (see Discussion).

We finally asked how the geometry of the representation relates to neuronal physiology. Are neurons essentially exchangeable as coordinate axes of the high-dimensional sensory representation or do different neuronal types participate in a distinct manner? One of the foremost physiological distinction made between cortical neurons is whether they are excitatory or inhibitory thus it is natural to ask whether they take on distinct roles in population geometry. In the three recordings where GABAergic neurons were identified with a tdtomato label we found systematic difference in the eigenmode loading’s on these putative GABAergic inhibitory neurons. For example in the second eigenmode of an example recording there is a large difference in the distribution of inhibitory neurons with positive eigenmode loadings (75 %) whereas other neurons in the recording are equally likely to have negative or positive loadings (Fig 5C). This provides evidence that the features encoded by the eigenmode have a distinctly more uniform effect on the activity of inhibitory interneurons than other neuron types. There was often a significant difference in the fraction of negative loadings between inhibitory neurons and other neurons across the top twenty modes (Fig 5D, green dots). Thus we find evidence that the principal stimulus features driving population responses have a distinct effect on inhibitory neurons. Specifically inhibitory neurons tuning includes a component of one sign of eigenmode tuning more often than other neurons. Understanding this tuning and the significance of one sign vs the other could be relevant to the function of inhibitory neurons in shaping sensory representations (see Discussion).

## Discussion

### Summary

We have introduced a novel and highly accurate estimator of the eigenspectrum of high-dimensional population neural tuning. In particular it performs well in the challenging conditions of limited stimuli and correlated noisy measurements that are common in large scale neural recordings. We applied this estimator to re-analyze a large scale recording of mouse primary visual cortex in response to natural images. We showed that the eigenspectrum was not well fit by a power law—in contrast to the conclusions prior work. Instead it was captured by a broken power law. The broken power law showed a characteristic form with an initially shallow slope for the first 10 eigenvalues (*α*1 ≈0.5) and a steeper fall off for the remaining eigenvalues (*α*2 ≈1.2). The tail of the signal eigenspectrum was steeper than previously estimated, *α* ≈ 1 vs *α* ≈1.2. We examined the image features that drove the dominant variation in the initial component of the power law and found their encoding fidelity was higher than the average neuron and that they sometimes were well characterized by classic models of primary visual cortex but also sometimes decidedly not. Finally we found that the features driving the dominant eigenmodes had distinct effects on putative inhibitory neurons, tending to be uniform in the sign of its effect. We thus have discovered clear links between geometry, computation and physiology in mouse primary visual cortex and introduced a novel estimator of high dimensional geometry that is more accurate than prior methods.

### Relevance to prior work

We re-analyzed the data of Stringer et. al., (3) and came to qualitatively different conclusions about the form of the signal eigenspectrum. Specifically, it was claimed that the signal eigenspectrum follows a power law with a slope near one whereas we found the signal eigen-spectrum is consistent with a broken-power law with neither of its slopes near 1. The authors originally argued that a slower-decaying eigenspectrum indicated a more efficient representation, whereas steeper decay reflected a smoother representation and that a power law with a slope of one was the slowest the eigenspectrum could decay (for the purpose of efficiency) before the representation became pathologically unsmooth. Thus their original cvPCA based estimates that the slope of the tail of the eigenspectrum was near 1 indicated that these theoretical considerations could precisely predict an empirical property of primary visual cortex. Yet, our more accurate MEME estimator revealed the slope was not at this critical point, weakening the explanatory power of their theory.

One explanation for deviations from their theory could rest in the veracity of its assumptions. The truth of the claim that a more slowly decaying eigenspectrum is in general more efficient depends on the form of the noise both in the responses of the neurons and in the stimulus. For example Atick and Redlich (26) found that when noise in inputs was low then the most efficient linear sensory transformation would whiten the inputs, thus the output eigenspectrum would be flat, but when noise was high it would average over inputs and the output eigenspectrum would fall off steeply. Thus an explanation for the signal eigenspectrum in mouse primary visual cortex being consistent with a broken power law may derive from the character of input and neuronal noise and not require consideration of constraints on the smoothness of the neural code.

Thus a fundamental feature of sensory representation of primary visual cortex, the form of its eigenspectrum, remains unexplained. Despite this mystery, our empirical finding has concrete consequences to the project of characterizing primary visual cortex that we discuss below.

### Interpretation of the signal eigenspectrum

Our estimator allows accurate estimates of the entire signal eigenspectrum of neural populations. We now consider two interpretations of the signal eigenspectrum of practical significance to understanding sensory coding: (1) its relevance to predictive modeling of neural tuning and (2) sensory encoding.

The signal eigenspectrum of a population of sensory neurons quantifies the optimal performance of a linear combination of image features in predicting the responses of those neurons. The cumulative sum up to the *n*th eigenvalue is exactly how much variance can be explained by *n* of these hypothetical optimally predictive image features. This puts a tight upper bound on the performance of the now common practice of regressing learned image features on neural responses (e.g.,DNN responses). If there are *n* features the variance explained cannot surpass 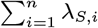. Thus an accurate estimate of the signal eigenspectrum could be used as a metric of how close a model is to optimal efficiency i.e., uses no more features than necessary for a given predictive performance. Thus a very practical view of the signal eigenspectrum is an exact quantification of minimal complexity of the model needed to capture the tuning of a population of neurons. A power law is a heavy tailed distribution which suggests the complexity of sensory representation is quite high—the performance of this hypothetical perfect model converges slowly with the number of parameters. Yet our finding of a steeper slope in the tail indicates substantial savings. For example to achieve 75% variance explained ∼900 features would be needed if the signal eigenspectrum followed a power law with a slope of 1 whereas for the broken power law on average ∼350 features are needed. In short, this study suggests that sensory neuroscientists seeking a compact but fairly predictive model of primary visual cortex should bear in mind they will need at least 350 image features—a large but still feasibly characterized number of features.

Alternatively, the image features of this hypothetical optimal model are also just image features that the neural population jointly encodes. Thus the eigenspectrum exactly quantifies the dimensionality of the features space within which the mean population neural response encodes images. From this perspective our finding of an initially slow decay of the eigenspectrum implies that there are ten or so roughly equally weighted features that the neural population encodes with high redundancy across a large population of neurons. The more rapid fall off in the tail of the signal eigenspectrum indicates that additional features quickly diminish in their effect on the population, but the heavy tail of the power law still insures that cumulatively these additional features drive the majority of variation (first 10 explain ∼30% of variation). While we have measured the degree of variation along these feature dimensions we do not know what these features are. Predictive models have primarily focused on the tuning of individual neurons, our measurements of signal eigenspectrum and associated SNR, indicate that the features encoded across the population are highly redundant and perhaps more relevant to downstream processes given their fidelity. Some of the variation in the tuning of individual neurons may reflect components that are not strongly represented at the population level (i.e., tuning that is unique to each neuron). Thus it could be productive to use predictive models to explain eigenmode tuning in addition to single neuron tuning. It may turn out that dominant modes are more easily captured similarly to how we found a surprising amount of their variation could be explained with a linear model.

### Population sensory representations in primary visual cortex

We found that the image features associated with the dominant eigenmodes were far more robustly encoded than those of the average neuron. This a clear empirical reason to recommend studying this population level tuning: primary visual cortex encodes these visual features in particular with very high fidelity. Further experiments where populations receptive fields are aligned to the same stimuli could determine whether this tuning is shared across animals.

Ultimately the striking difference in SNR between neurons and eigenmodes is the result of commonality in tuning across the population of neurons—signal correlation—where tuning redundancy leads to robustness to noise. While long hypothesized to be a potential consequence and normative explanation of signal correlation (21–23) direct estimation of what features are robustly encoded in primary visual cortex is enabled by simultaneous recordings of a large population of retintopically overlapping neurons. Stereotyped properties with respect to selectivity for spatial frequency and orientation in primary visual cortex has long been known but it has become increasingly clear that primary visual cortex responses are not solely characterize by their selectivity for spatial frequency and orientation (6, 7). Thus our finding that, similarly, eigenemodes are often not well-described by such classical notions is a data-driven indication that there are uncharacterized but stereotypical components of single neuron tuning encoded across the population. We have not exhaustively characterized these modes with respect to the more recent models of primary visual cortex (e.g., inclusion of a normalization pool (27)) the significance of these more recent efforts could be emphasized if they are shown to capture unexplained population level representation. Otherwise more flexible data driven models (e.g., deep neural networks) could be applied to eigenmode tuning and the difficult work of characterizing these models could in part be justified by the assurance they were capturing image features primary visual cortex robustly encodes at the population level.

### Inhibitory neurons distinct participation in sensory representation

Inhibitory neurons, in contrast to excitatory neurons, typically do not have axons that extend to other regions of visual cortex—thus they presumably act to modulate the sensory representation that is transmitted to other brain regions (28).

There is evidence that inhibitory neurons, when collectively activated, can sharpen tuning (8). We found that putative inhibitory neurons had the same sign eigenmode tuning more often than other neurons—in other words inhibitory neurons tended to co-activate at a higher rate than other neurons in response to features that maximally drove the population. Future work could causally test if inhibitory neurons sharpen tuning along directions of eigenmode tuning. This could reveal novel computational roles of inhibitory interneurons in shaping the visual sensory representation beyond single neuron orientation and spatial frequency tuning.

## Conclusion

We have made several principal empirical observations about the population code in primary visual cortex. The signal eigen-spectrum is a broken power law with slow than rapid decay, the top modes are of far higher fidelity than the average individual neuron and are often not well characterized by classic models of primary visual cortex, and inhibitory neurons tend to be driven in concert by the top modes’ features. Taken together these results challenge the primacy of studying individual neural tuning curves given the dramatic emergence of distinct, but poorly characterized, population level computations that are robust to noise, and with clear relevance to physiology.

Beyond our empirical findings, we have demonstrated that the challenge of describing high dimensional neural codes requires novel statistical methods that are rigorously validated. They lay a critical foundation for surmounting the ‘curse of dimensionality’ in the study of neural representations and motivate addressing this curse because they indicate the rich statistical structure that lays waiting to be uncovered at the population level.

## Materials and Methods

### Assumptions and terminology for derivation of estimator

Here we employ a common model of population neural responses:

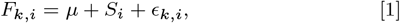

where *F*_*k,i*_ is a vector of responses from *n* neurons to the *k*’th repeat of the *i*’th stimulus, *µ* is a vector of the mean (across the stimuli distribution) responses of each neuron, *S*_*i*_ is the vector of expected neural responses to the *i*th stimulus, (i.e., samples from the tuning curve) with signal covariance Σ_*S*_, and *E*_*k,i*_ is the per-trial noise with noise covariance Σ_*N*_. The signal covariance Σ_*S*_, the object of our current study, is given by the covariance of noiseless responses *S*_*i*_ over the stimulus distribution *P* (*S*) (e.g., sampling from a database of natural images). We will often deal with *m × n* matrices of responses collected on the *k*th repeat which we will call *F*_*k*_, the concatenation of *m* draws from *F*_*k,i*_. Here we focus on the estimation of the signal eigenvalues, *λ*_*S,i*_ = *f*_*i*_(Σ_*S*_), the sorted eigenvalues of the signal covariance matrix Σ_*S*_. We also consider the noise eigenvalues, *λ*_*N,i*_ the sorted eigenvalues of the noise covariance matrix Σ_*N*_. We will estimate these quantities indirectly from unbiased estimates of signal and noise eigenmoments, the *p*’th moments respectively being: 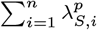 and 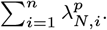.

### Unbiased estimation of eigenmoments

Our estimator infers the signal eigenspectrum by matching unbiased estimates of signal eigenmoments. It is an extension of previous work developing unbiased estimates of eigenmoments from noiseless data that we review next.

### Estimation from noiseless data

Unbiased estimates of eigenmoments were first discovered by Li *et al* (29) but then employed to infer eigenspectra by Kong and Valiant (30). In the noiseless case we have direct observations of *S*_*i*_ (letting *µ* = 0), and an *m × n* matrix formed by concatenating *m* presentations of stimuli, we call *S*. For insight into the method we show how an unbiased estimate of the *p*th eigenmoment can be calculated in the noiseless case. A single unbiased estimate of covariance can be formed from the *j*th observation 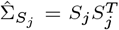 then the statistic 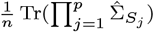, formed from *p* independent estimates of covariance, is an unbiased estimate of the *p*th eigenmoment because,

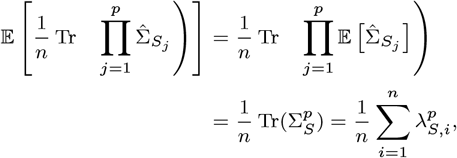

the first step following from independence of the estimates of covariance and the linearity of the trace and expectation, the second step is by definition true of an unbiased estimate, and the last step follows from the identity for symmetric matrices 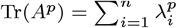.

It is unnecessary to explicitly calculate the outer product for each 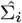 but instead calculate inner products, for example,

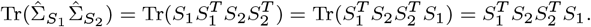

More generally we can get the *p*th eigenmoment as follows, let *σ* be a set of *p* distinct indices of IID observations of *S* [*σ*_1_, *σ*_2_, …*σ*_*p*_] and *σ*_*p*+1_ = *σ*_1_. Then the estimator of the *p*th eigenmoment is,

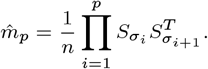

As *p* and *m* grow there are many number of distinct indices over which to form this estimator. To reduce variance one could average over all possible sets of distinct indices. This quickly becomes computationally intractable and so Kong and Valiant developed a rapid approximation where they average over all increasing sets of indices. This can be accomplished with the following calculation letting *A* = *SS*^*T*^, where *S* is the *m × n* concatenation of *m* observations and *A*up be the same matrix with lower triangular and diagonal entries set to 0,

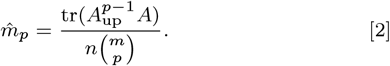

### Extension to noisy data

The estimator of Kong and Valiant assumed that there was no noise, correlated or otherwise, in the measurement of *S*. Here we extend their estimator to the case of measurement error as described in Eqn.1. The key insight is that across repeats noise, *E*_*k,i*_, will be independent while signal, *S*_*i*_, will be identical. So, under the assumption *µ* = 0, we obtain the *i*th unbiased estimate of the signal covariance with only two repeats of data 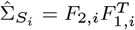 because,

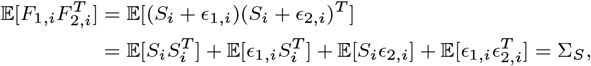

where the last step follows from the independence between signal, noise, and different trials of noise. Following the logic of the prior section we can set 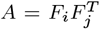 where *i* ≠ *j* and Eqn. 2 serves as our unbiased estimator. Importantly the unbiased nature of this estimator does not depend on the distribution from which data are drawn. For the common case where *µ* ≠ 0 we transform our data so that *µ* ≠ 0 but the covariance remains the same, then apply our estimator to this transformed data. The transformation is simple, for each repeat separate the responses into two disjoint sets of stimuli responses (same number of stimuli in each) take their difference and divide by 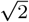. This works because draws of noise and signal across stimuli are independent but the mean response is constant, and so the difference only scales signal and noise covariance while removing the mean. In our analyses, we take the strategy of subtracting the odd stimuli from the even stimuli for each repeat. The variance of this procedure could be reduced by taking more differences from the many possible disjoint sets and calculating estimates of eigenmoments for all of them and then averaging. Naively, we could have subtracted the sample mean from each neuron calculated across all stimuli. This would change the covariance structure of the observations resulting in an unnecessary bias, thus we do not take this approach.

### Fitting the eigenspectrum using estimated eigenmoments

The goal of the methodological developments in this paper is to infer eigenspectra from finite noisy data. Kong and Valiant develop a non-parametric approach to this inference in the noiseless case. Here we propose a parametric approach for the cases where scientific questions pertain directly to parameteric forms of eigenspectra (e.g., Stringer *et al*, (3)). In addition, if a parametric form can be assumed then there are potentially large gains in accuracy to be made. We can also assess if the parametric form is a good assumption by determining if it can account for systematic variation in the eigenmoments (see below).

Our approach to inferring parametric eigenspectra is simple: we optimize the parameters of the assumed form of the eigenspectrum to minimize the squared error between its eigenmoments and the eigenmoments estimated from the data. To solve the nonlinear least squares problem and satisfy constraints on parameters (e.g., power law slope cannot be negative because eigenvalues monotonically decrease) we use the nonlinear least squares function implemented in scipy (31). When fitting a broken power law we simply perform a grid search of potential breaks points, optimize slope and intercept parameters for each, then use the break points that gave the minimal error. In practice we scale the variance of the data before estimating the eigenmoments because later eigenmoments can easily go beyond the floating point range if the raw scale of the data is too high or low. We scale the data with an unbiased estimate of the total signal variance (sum of signal variance across all neurons). The eigenspectrum of the raw data can easily be recovered by re-scaling. Estimated eigenmoments are heteroscedastic and correlated which can affect the accuracy of this estimation procedure. Higher-order eigenmoments tend to be increasingly variable thus including them can make estimates of eigenspectrum parameters unstable. To address this we estimate the sampling covariance matrix of the estimated eigenmoments with a bootstrap procedure (sample with replacement from stimuli) and then apply a whitening matrix to the errors between the estimated and fit eigenmoments. This effectively weights the eigenmoments according to their reliability. In practice we find that parameters are not changed by using more than 10 eigenmoments because their variability is extremely high and thus their influence is down weighted.

### Simulation procedure

To validate our estimator and create parametric bootstrap confidence intervals around our estimates we make use of simulations that match the distribution of the original experimental data. We simulate according to the model specified in Eqn. 1. We form an unbiased estimate of the noise covariance for each stimulus by subtracting off the mean of the two trials than averaging these individually unbiased estimates across all stimuli as follows,

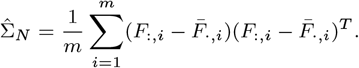

To form an unbiased estimate of the signal covariance we calculate the sample covariance between the two repeated observations of stimuli,

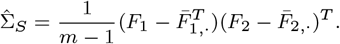

Neither estimate will be positive semi-definite (PSD) because there are fewer stimuli than neurons. Furthermore the estimate of signal covariance is unlikely to be symmetric. To address this we force the noise covariance matrix to be PSD by finding its eigenvalues and setting any less than 0 to be 0. To force the signal covariance matrix to be symmetric we average it with its transpose, then force this covariance matrix to be PSD.

In most simulations we set the signal eigenspectrum by performing the eigenvalue decomposition, 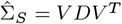, then reconstructing the covariance matrix but with the desired eigenvalues, *D*, giving 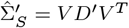.

### cvPCA

We calculate the cvPCA estimator for the *i*th signal eigenvalue,(*λ*_*S,i*_) as follows,

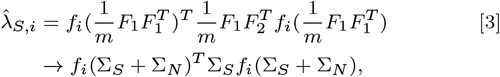

where *f*_*i*_() calculates the *i*th eigenvector via SVD, the arrow indicates convergence as *m* → ∞, and we assume *µ* = 0 (see Eqn. 1). Thus it estimates an eigenvector from the covariance estimated in one repeat of data and then finds the amount of variance explained by it in an unbiased estimate of the signal covariance calculated across different repeats. This is not a consistent estimator because the eigenvector estimates converge to those of Σ_*S*_ + Σ_*N*_, thus depending on the relationship between signal and noise these estimates can be inaccurate. For example if they have the same eigenvectors in the same ordering then they will converge to the correct value (except for those beyond the rank of the data), whereas if they are independent this step of cvPCA will mis-estimate the directions to calculate maximal variation in the unbiased estimate of signal covariance. The calculation method in Stringer *et al* 2019 (3) differs slightly from that here, the principal difference being that singular vectors are calculated directly from neural responses instead of their sample covariance. We confirmed that this approach gives essentially the same numerical results while having a simple form from which the estimator’s inconsistency is clear.

### Consistent estimates of eigenmode tuning and loadings

As shown above using raw data to estimate signal eigenvectors can lead to gross biases. Here we analyzed the tuning of signal eigenmodes and the loadings on individual neurons. Thus we sought a consistent estimator of these quantities. We estimated these respectively by performing SVD on 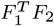, the unbiased estimate of signal covariance between stimuli, and 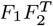, the unbiased estimate of signal covariance between neurons. As long as the signal eigenvalues are monotonically decreasing this provides consistent estimates of eigenvectors associated with signal eigenvalues above the rank of the data.

### Estimation of model performance

We sought to evaluate the fraction of signal variance explained by our models. Applying the naive estimate of *R*^2^ between neurons and a model’s prediction would be downwardly biased by trial-to-trial variability and upwardly biased by the number of model parameters—over-fitting (24). Thus we estimated model performance with a noise and model degrees of freedom corrected estimator. Below we provide a short derivation. We rewrite the model of population neural responses by separating *S*_*i*_ into two terms,

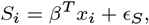

where *x*_*i*_ with *Cov*[*x*_*i*_] = Σ_*s*_ is the vector of *d* model feature values in the *i*th image, *β* is the fixed set of weights that determines the linear relationship between the neural signal and image features, and *E*_*S*_ is the component of the neural signal that cannot be predicted by a linear combination of the model features that is assumed to be distributed as *E*_*S*_ ∼ *N* (0, *σ*^2^).

Our desired estimand is then,

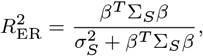

which goes to 1 if the neural signal responses are a perfect linear function of the features *x*_*i*_. To estimate this quantity we follow the approach of finding unbiased estimates of the numerator and denominator.

For the numerator, under the above assumptions, the residual sum of squares from the least square fit of image feature to neural responses is distributed as follows,

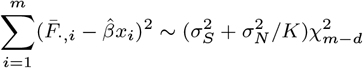

So,

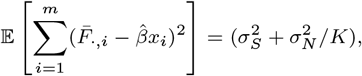

an unbiased estimate of total variance is,

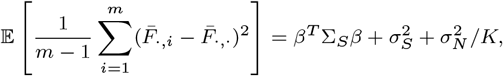

so subtracting the two estimators gives an unbiased estimate of linear variance,

For the denominator an unbiased estimate of trial-to-trial variability can be subtracted from the unbiased estimate of total variance.

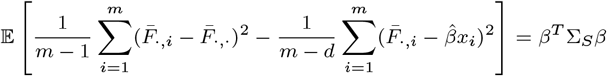

To estimate linearity of tuning we regressed the pixels of images on response profiles. The raw stimuli are grey scale images of 68 × 270 pixels giving 18, 360 features in the regression but there were only ∼2, 800 stimuli shown thus the linearity statistic cannot be naively calculated (i.e., there are more features than observations). To address this we performed principal components regression: we regressed onto the top *D* principal components of the images. We found that performance saturated at *D* ≈ 100 principal components of the images, we used *D* = 100 features for the *R*^2^ values in Fig 4B and 5B.

To determine if a population of class simple and complex cell models, could account for additional variance we formed a basis of gabor filters tiling scale (fraction of image 1, 0.5, 0.25), orientation (4 rotations), phase (0 and 90 degrees), and position (non-overlapping tiling at each scale). We then regressed on the responses of these filters and their square to calculate *R*^2^ as described above. To tile the entire 68 × 270 image with gabor filters would result in more features than observations. Fortunately we found that it was unnecessary to use the entire image because receptive fields in each recording were restricted to a small subset of the image. To localize receptive fields we estimated the linear receptive fields of all neurons in each recording we then took the sum of squares across neurons as an estimate of the population receptive field profile. We estimated the boundary of the receptive field with the 95th quantile of its profile and extracted a rectangular patch of the image that contained the boundary of the receptive field (for example, Fig 3D red dashed box). We found that the performance of a linear model predicting eigenmode responses did not significantly suffer when using this restricted patch instead of the entire image. We also used these receptive field bounds to visualize linear receptive fields (Fig 4C,F).

### Experimental data

All stimuli were presented for 0.5s with a random inter-stimulus interval between 0.3 and 1.1s consisting of a grey-screen. The images used in the experiment were taken from the ImageNet database, which includes categories such as birds, cats, and insects. The researchers manually selected images that had a mix of low and high spatial frequencies and that did not consist of more than 50 % uniform background. All images were uniformly contrast-normalized by subtracting the local mean brightness and dividing by the local mean contrast. To compute the local mean and standard deviation, a Gaussian filter with a standard deviation of 30 degrees was used. Each stimulus consisted of a different normalized image from the ImageNet database, with ∼2, 800 different images used in total. The same image was displayed on all three screens, but each screen showed the image at a different rotation.

Mice bred to express GCaMP6 in excitatory neurons were used in the majority of recordings. Mice bred to express tdTomato in inhibitory neurons were also used in a subset of the recording while GCaMP6 was expressed virally, allowing the identification of inhibitory and excitatory neurons.

Neural activity was recorded using a two-photon microscope while the mice were free to run on an air-floating ball. Recordings were collected across multiple depth planes at a frequency of 2.5 or 3 Hz, with planes 30-35 *µm* apart. The field of view of the microscope was selected such that 10,000 neurons could be observed with a retinotopic location on the stimulus display. The 2,800 natural image stimuli were displayed twice in a recording in two blocks of the same randomized order.

Calcium movie data was processed using the Suite2p toolbox to estimate spike rates of neurons. Underlying neural activity was estimated using non-negative spike deconvolution. These deconvolved traces were normalized to the mean and standard deviation of their activity during a 30-minute period of grey-screen spontaneous activity. For further detail please see the original study. All analyses done in this paper were performed on the pre-processed data available on figshare (32) (https://figshare.com/articles/Recordings_of_ten_thousand_neurons_in_visual_cortex_in_response_to_2_800_natural_images/6845348).

## ACKNOWLEDGMENTS

We thank members of the Pillow lab, Kenneth Harris, Matteo Carandini, Carsen Stringer, and members of the Cortexlab for for their feedback on the project.

## Supporting Information Appendix (SI)

**Fig S1.**
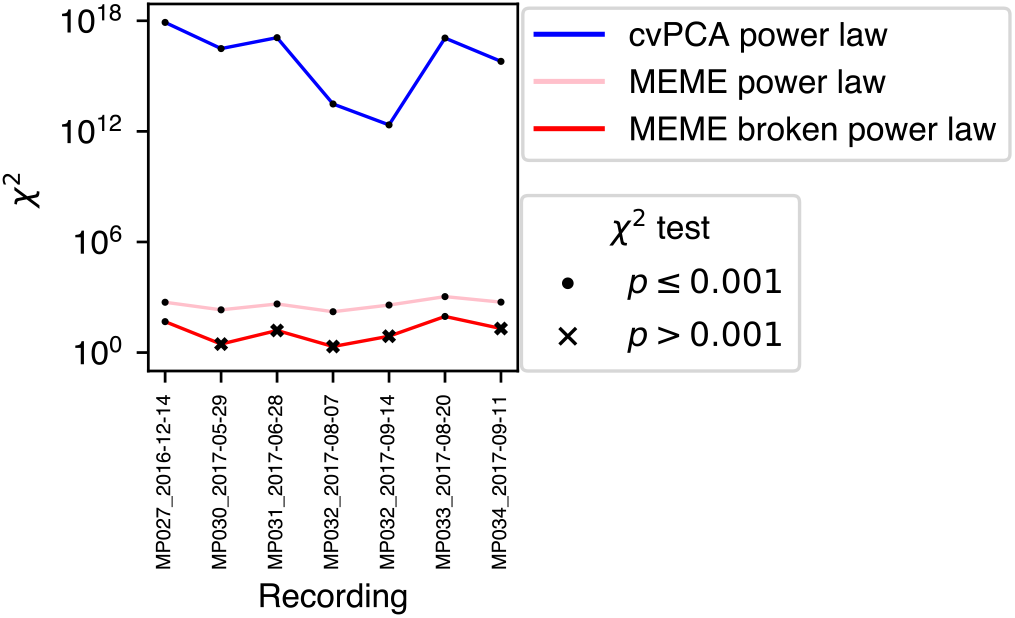
Chi-squared test statistic of difference between eigenmoments of parametric model of eigenspectrum and direct unbiased estimates of eigenmoments from data. This was the sum of squared weighted errors (see Methods, Fitting the eigenspectrum using estimated eigenmoments) and the null distribution was Chi-squared with degrees of freedom (DOF) equal to the number of model parameters (cvPCA DOF=2, MEME power-law DOF=2, MEME broken power-law DOF=4).

**Fig S2.**
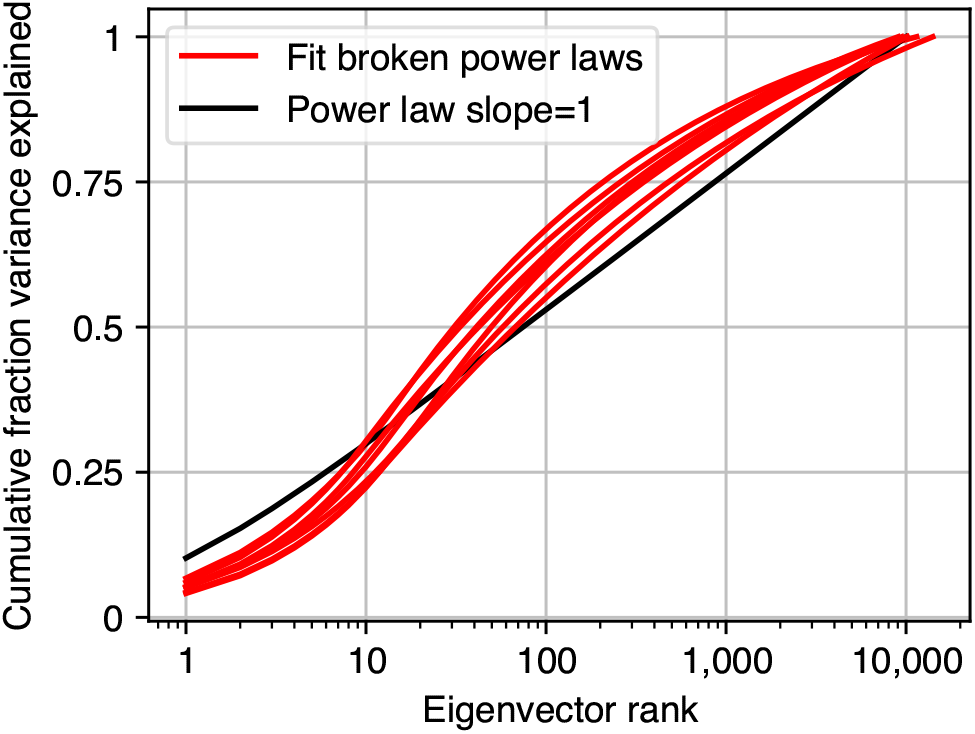
Cumulative fraction signal variance explained as a function of eigenvector rank. Broken power-laws fit to 7 recordings of responses to natural images are plotted in red and for reference a power law with slope of 1 is plotted in black (10,000 neurons).

**Fig S3.**
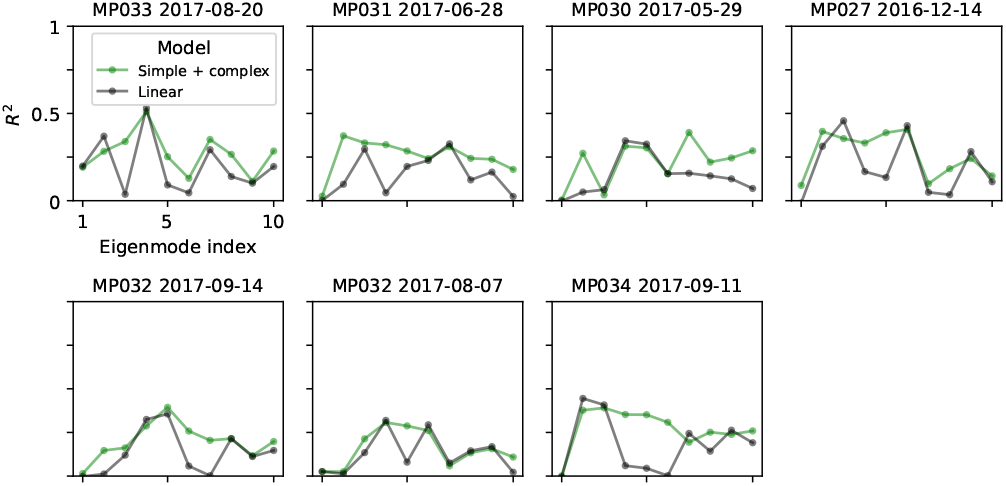
For all recordings the fraction variance explained (corrected for model degrees of freedom) of the top ten eigenmodes by a linear model (black) and a simple and complex cell model (green). See Methods, Estimating model performance.

## Notes

https://doi.org/10.25378/janelia.6845348.v4

### Competing Interest Statement

The authors have declared no competing interest.

